# Recursive Cleaning for Large-scale Protein Data via Multimodal Learning

**DOI:** 10.1101/2024.10.08.617190

**Authors:** Zixuan Jiang, Sitao Zhang, Jiahang Cao, Qiang Zhang, Shiyi Liu, Yuetong Fang, Lingfeng Zhang, Rui Qing, Renjing Xu

**Affiliations:** Hong Kong University of Science and Technology (Guangzhou); Shanghai Jiao Tong University

## Abstract

Reliable datasets and high-performance models work together to drive significant advancements in protein representation learning in the era of Artificial Intelligence. The size of protein models and datasets has grown exponentially in recent years. However, the quality of protein knowledge and model training has suffered from the lack of accurate and efficient data annotation and cleaning methods. To address this challenge, we introduce **ProtAC**, which corrects large **Prot**ein datasets with a scalable **A**utomatic **C**leaning framework that leverages both sequence and functional information through multimodal learning. To fulfill data cleaning, we propose the Sequence-Annotation Matching (SAM) module in the model, which filters the functional annotations that are more suitable for the corresponding sequences. Our approach is a cyclic process consisting of three stages: first pretraining the model on a large noisy dataset, then finetuning the model on a small manually annotated dataset, and finally cleaning the noisy dataset using the finetuned model. Through multiple rounds of “train-finetune-clean” cycles, we observe progressive improvement in protein function prediction and sequenceannotation matching. As a result, we achieve **(1)** a state-of-the-art (SOTA) model that outperforms competitors with fewer than 100M parameters, evaluated on multiple function-related downstream tasks, and **(2)** a cleaned UniRef50 dataset containing ∼50M proteins with well-annotated functions. Performing extensive biological analysis on a cleaned protein dataset, we demonstrate that our model is able to understand the relationships between different functional annotations in proteins and that proposed functional annotation revisions are reasonable.

## 1 Introduction

Proteins, central components of cellular machinery, have been the focus of extensive experimental and computational approaches aimed at elucidating their functions. The advent of high-throughput sequencing technologies Reuter et al. (2015) has led to a significant increase in the number of sequenced genomes in past two decades, resulting in the creation of extensive protein databases. These databases serve as training resources for the advancement of deep learning in protein research Chen et al. (2024); Elnaggar et al. (2021); Lin et al. (2023); Ferruz et al. (2022); Nijkamp et al. (2023).

Language models (LM) are highly valued for their effectiveness in natural language processing, and a specialized version known as Protein Language Model (PLM) has been extensively utilized in protein representation learning. This variant leverages protein sequences as training data, as amino acid sequences serve as the fundamental coding for proteins. PLMs demonstrate exceptional capabilities in comprehending protein functions Rives et al. (2021); Brandes et al. (2022); Meier et al. (2021); Vig et al. (2020) and structures Rives et al. (2021); Lin et al. (2023); Rao et al. (2020); Vig et al. (2020), thereby facilitating *de novo* protein design Verkuil et al. (2022).

Recent studies Xu et al. (2023); Zhang et al. (2023) have demonstrated that PLMs leveraging multimodal information, such as sequence, functional annotation, and structure data from proteins, exhibit superior capabilities compared to models pretrained solely on sequences. However, despite significant advancements in computational methods, particularly in deep learning, which have achieved near laboratory-level precision in protein structure prediction Jumper et al. (2021); Baek et al. (2021); Abramson et al. (2024) and expanded the structural coverage of the known protein-sequence space Varadi et al. (2022), accurate protein function prediction remains a challenge. High-quality protein structure databases Burley et al. (2017) and biological knowledgebases Boutet et al. (2007) are still relatively limited in scale compared to the vast amount of validated sequences available. Existing automatic annotation methods, primarily statistical and rule-mining-based approaches Consortium (2019) applied to largescale protein datasets, often face challenges when applied to large-scale protein datasets due to the complex mapping between protein sequences and functions, resulting in inaccuracies in protein property annotations. These issues not only impact data quality but also introduce uncertainty into subsequent research endeavors. Therefore, the identification and removal of noise and errors to enhance the accuracy and reliability of protein datasets are crucial in the fields of bioinformatics and proteomics. Effective solutions are urgently needed to address these challenges.

Building upon the latest advancements in protein multimodal learning methods Xu et al. (2023); Brandes et al. (2022), we propose an innovative learning framework that integrates multiple modalities of protein data, including sequence and functional information. Drawing inspiration from the concept of matching in Vision-Language Learning Li et al. (2021; 2022), our framework effectively discerns between reliable and unreliable information within large-scale protein datasets. Our approach introduces a novel multi-round training strategy, where each round involves model pretraining on a noisy dataset followed by finetuning on a manually curated dataset. Subsequently, the model is tasked with cleansing the noisy dataset by predicting and selecting credible protein function information. The cleaned dataset is then recursively utilized in the next round of pretraining. A visual representation of the concept of recursive data cleaning for protein datasets is depicted in Fig.1a. This iterative process enables the replacement of noisy datasets in subsequent rounds, leading to mutual enhancement of both dataset quality and model performance.

**Figure 1:**
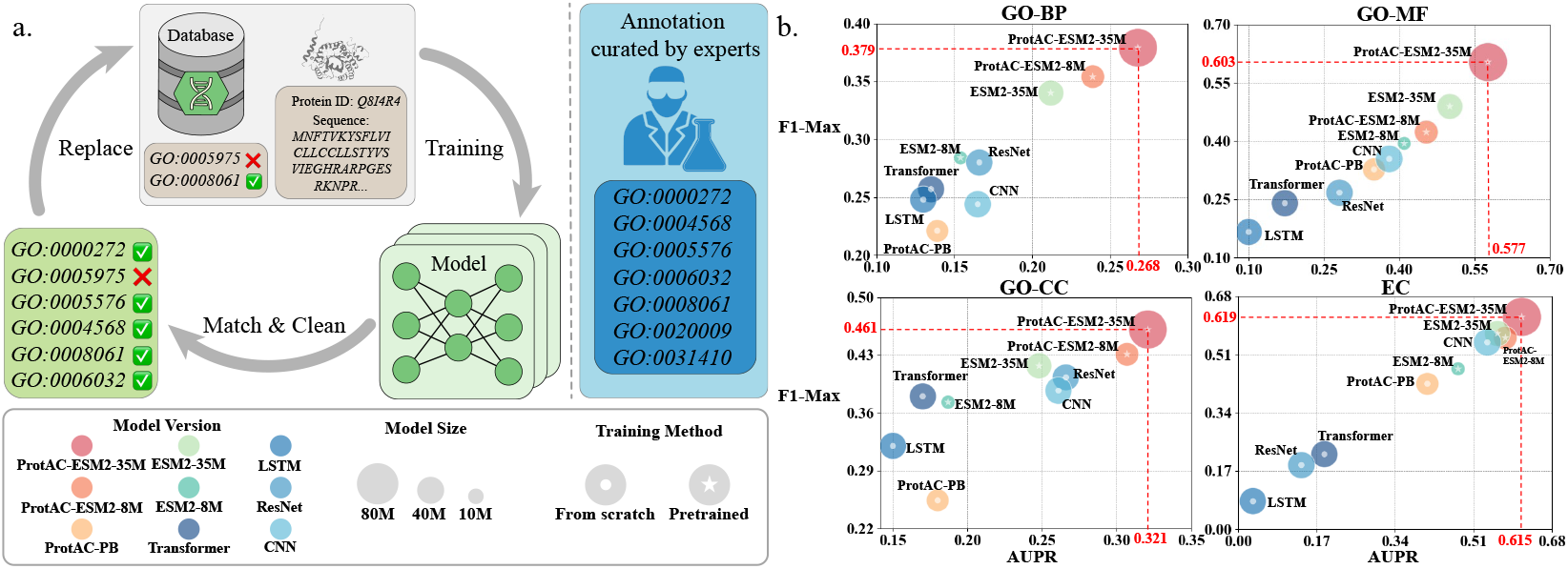
**(a)** Schematic diagram of recursive data cleaning for protein function annotation and expert-curated ground-truth annotations. We take protein ID Q8I4R4 as an example. This cycle is repeated many times, and the modified annotations are more consistent with the results of manual screening by biologists than the original annotations in the database. **(b)** Performance of ProtAC and other models with less than 100M parameters on downstream tasks related to function prediction.

Our study explores efficient model training approaches and finds that using pretrained weights is more effective than training from scratch for enhancing model training and dataset quality. Models with larger parameter sizes outperform smaller models in functional prediction tasks and dataset quality improvement. Our model achieves SOTA results in protein function prediction tasks, surpassing models with similar parameter sizes (Fig.1b) and remaining competitive against larger PLMs. We evaluate the data cleaning capabilities of our model using the newly updated SwissProt dataset, which experts have validated but the model has never seen. This evaluation shows that our model has exceptional anomaly detection capabilities and greatly improves the accuracy of protein function prediction, nearly matching the proficiency of human biologists. Furthermore, we conduct a detailed biological analysis of the cleaned dataset, successfully verifying that our modifications to noisy protein information are biologically meaningful and align with biological principles.

## 2 Related Work

### Protein Multimodal Learning

Mutual understanding of sequence and function plays a significant role in exploring biological behaviors. Recently, multimodal models have been developed to integrate information from protein sequence and function. ProteinBERT Brandes et al. (2022) adopts the classical BERT architecture and leverages local attention to integrate protein sequence information and utilizes global attention to learn function information; OntoProtein Zhang et al. (2022) learns protein representations under the context of a knowledge graph; ProGen Madani et al. (2020) incorporates protein function labels to generate functional proteins. However, none of the above models considers the role that biomedical text can play. ProtST Xu et al. (2023) takes the initiative of enhancing both representation learned by protein sequence and biomedical texts.

### Protein Functional Annotation Prediction

The accuracy of protein function prediction is an important reflection of PLM capabilities. Gene Ontology (GO) Ashburner et al. (2000) annotations provide a detailed description of protein functions in biological systems. Predicting GO annotations for uncharacterized proteins is crucial for exploring unknown protein landscapes. In each Critical Assessment of Functional Annotation (CAFA), several noteworthy protein GO annotation prediction models appear Yao et al. (2021); Wang et al. (2023); Kulmanov et al. (2018); Zhou et al. (2019), showing significant progress. AnnoPRO Zheng et al. (2023) combines protein sequence representation with GO functional family information to capture the intrinsic correlation between protein features and significantly improve the annotation performance of low-abundance protein families.

### Knowledge Distillation and Data Cleaning

Knowledge Distillation (KD) Hinton et al. (2015) aims to improve the performance of student models by distilling knowledge from teacher models. Different from most existing KD methods, which simply force the student to have the same category predictions as the teacher, Li et al. (2022) proposed CapFilt, which can be interpreted as a more effective way to perform KD in the context of Vision-Language Pretraining (VLP), where the captioner distills knowledge through semantically rich synthetic captions, while the filter distills knowledge by removing noisy captions. We apply this idea to large protein dataset cleaning for the first time. See more related work in Appendix A.

## 3 Method

We present ProtAC, an automatic data cleaning framework for large protein datasets that leverages unified knowledge from protein sequence and functional annotations. In this section, we first introduce our model architecture and its training objectives, and then describe our data cleaning strategy.

### 3.1 Model Architecture

#### Overview

Our model structure consists of four main components: Sequence Encoder, Annotation Encoder, Annotation Decoder, and SAM Filter (the model architecture is shown in Fig.2). As a versatile learning framework, the first three main modules can be easily replaced with mainstream PLMs, which greatly expands the scope of further model design and lays the foundation for inspiring future directions of protein multimodal representation learning.

**Figure 2:**
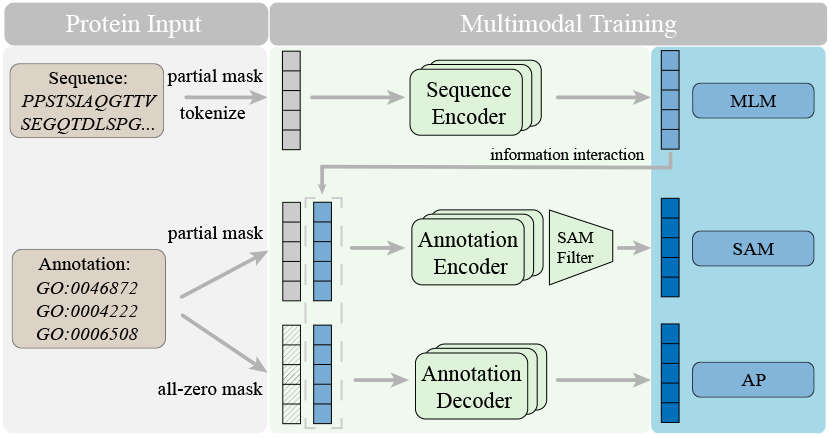
Model architecture and training objectives of ProtAC.

#### Sequence Encoder

We investigate two widely used PLMs: ESM2 Lin et al. (2023), one of the current SOTA sequence-only PLMs, known for its superior protein feature extraction capabilities, making it a common choice for protein representation learning, especially in multimodal learning tasks; ProteinBERT Brandes et al. (2022), a multimodal PLM based on the BERT architecture, which captures sequence information through its local part and functional information through its global part, with a cross-attention mechanism promoting the interaction between the two parts. For our Sequence Encoder, we use the local part of ProteinBERT and ESM2. Input sequence is partially masked and tokenized. Considering the limitations of computing resources and time, we selected the 8M and 35M variants from the ESM2 family, which stand out among PLMs with less than 100M parameters, have excellent performance and are sufficient for our data cleaning task requirements. We consider introducing PLMs with billions of parameters in future research to achieve larger-scale and more reliable functional correction.

#### Annotation Encoder and Decoder

We modify the global part of ProteinBERT as our Annotation Encoder. The input annotation information in the form of a fixed-size binary vector is partially masked and encoded into the annotation embedding. The sequence information is injected by inserting an additional cross-attention layer between the feed-forward networks of each block of the Annotation Encoder. The output embedding is used as a multimodal representation of the sequenceannotation pair. Annotation Decoder adopts the same structure as Annotation Encoder. The input annotations (all masked to zero vectors) are combined with the sequence information injected through the cross-attention layer to predict the correct annotation list.

#### Sequence-Annotation Matching (SAM) Filter

This module consists of a simple linear layer that can identify whether the input sequence and annotation match by processing the fused features of the sequence-annotation pair. The output of the SAM filter [*P*_*unmatch*_, *P*_*match*_] is a two-dimensional vector where the two dimensions represent the probability of a match and the probability of a mismatch of the input sequence and annotation pair, respectively.

### 3.2 Upstream Tasks AND Objectives

#### Overview

We jointly optimize three objectives during training, one understanding-based objective and two generation-based objectives, and compute three losses to activate different modules, as shown below.

**Masked Language Modeling (MLM)** activates the Sequence Encoder. It aims to predict the identity of amino acids that have been randomly masked out of protein sequences:

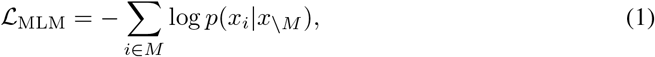

where for a randomly generated mask *M* that includes 15% of positions *i* in the sequence *x*, the model is tasked with predicting the identity of the amino acids *x*_*i*_ in the mask from the surrounding context *x*_*\M*_, excluding the masked positions. This masked language modeling objective Devlin et al. (2018) causes the model to learn dependencies between the amino acids. Although the training objective itself is simple and unsupervised, solving it over millions of evolutionarily diverse protein sequences requires the model to internalize sequence patterns across evolution.

**Sequence-Annotation Matching (SAM)** activates the Annotation Encoder. It aims to predict whether a pair of sequence and annotation is positive (matched) or negative (unmatched). Let *S* denote input sequence and *A* denote input annotation. We use Annotation Encoder’s output embedding as the joint representation of the sequence-annotation pair, and append the SAM Filter to predict a two-class probability *p*^*sam*^. The SAM loss is defined as the cross-entropy *H* between *p* and *y*:

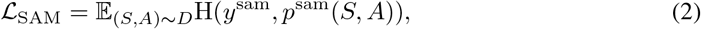

where *y*^*sam*^ is a 2-dimensional one-hot vector representing the ground-truth label. We follow the strategy proposed in ALBEF Li et al. (2021) to sample hard negatives for the SAM task with zero computational overhead. For each sequence in a mini-batch, we sample one negative annotation embedding from the same batch. Likewise, we also sample one hard negative sequence for each annotation. This results in the quantity of negative pairs being twice that of positive pairs for each mini-batch. Consequently, in practical training, we employ focal loss Lin et al. (2017) in lieu of cross-entropy loss to mitigate the adverse effects arising from the imbalance in sample quantities.

**Annotation Prediction (AP)** activates the Annotation Decoder. This loss minimized by Annotation Decoder during training is a sum of the categorical cross-entropy over the protein sequences and the binary cross-entropy over the annotations, namely

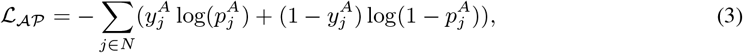

where *N* denotes the dictionary size of annotation list, 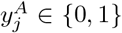 is the true label for annotation *j*, and 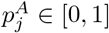 is the predicted probability that the protein has annotation *j*.

The **overall training objective** of ProtAC is:

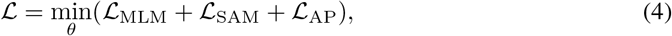

where *θ* denotes all trainable parameters including those of the three major modules and all projection heads. We seek to minimize the loss functions of all upstream tasks simultaneously during the training process.

### 3.3 Data Cleaning Workflow

#### Overview

We propose a multi-round data cleaning strategy with three stages in each round, namely pretrain, finetune and caption. Our core aim is to facilitate a reciprocal enhancement of model performance and dataset quality through a cyclical process, wherein the initialized model goes through the pretraining stage and the finetuning stage, producing a pretrained model and a finetuned model, respectively. Both models are subject to evaluation covering upstream and downstream tasks. The pretrained model will replace the initialized model in subsequent cycles, while the finetuned model will enter the caption stage to perform data cleaning on the noisy dataset, thereby producing a cleaned dataset that will replace the noisy dataset in the next cycle (data cleaning workflow is shown in Fig.3).

**Figure 3:**
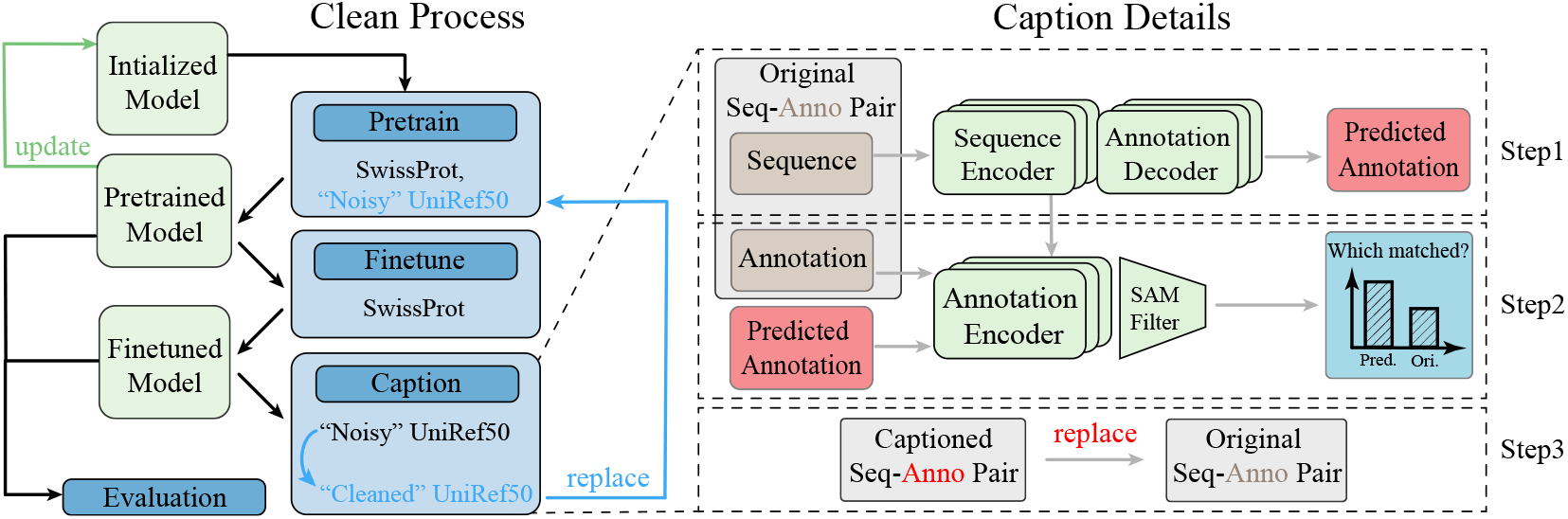
Data cleaning workflow of ProtAC. The left part of the figure outlines the cleaning process, and the right half of the figure details the caption process.

#### Stage Pretrain

Our models will load last-round pretrained weights except in the first round where methods for initializing models vary across different versions. Pretraining dataset is the combination of Uniref50 (noisy dataset) and SwissProt-trainset (well-annotated dataset).

#### Stage Finetune

Our model inherits the weights from Stage Pretrain and is finetuned for 10 epochs on SwissProt-trainset, which enables the model to have a high ability to distinguish between fake and real protein annotations, so the model utilizes the knowledge gained during finetuning and performs well in Stage Caption. The pretrained or finetuned models are evaluated by downstream tasks.

**Table 1:** Downstream task performance. We use three color scales of blue to denote the **first, second** and **third** best performance. *Abbr*., PB: ProteinBERT; Param.: Parameter.

| Pretrained Model | Param.<br><100M | GO-BP |  | GO-MF |  | GO-CC |  | EC |  |
| --- | --- | --- | --- | --- | --- | --- | --- | --- | --- |
| | | AUPR | $F_{max}$ | AUPR | $F_{max}$ | AUPR | $F_{max}$ | AUPR | $F_{max}$ |
| Naive Baselines |  |  |  |  |  |  |  |  |  |
| CNN Shanehsazzadeh et al. (2020) | ✓ | 0.165 | 0.244 | 0.380 | 0.354 | 0.261 | 0.387 | 0.540 | 0.545 |
| ResNet Rao et al. (2019) | ✓ | 0.166 | 0.280 | 0.281 | 0.267 | 0.266 | 0.403 | 0.137 | 0.187 |
| LSTM Rao et al. (2019) | ✓ | 0.130 | 0.248 | 0.100 | 0.166 | 0.150 | 0.320 | 0.032 | 0.082 |
| TransformerRao et al. (2019) | ✓ | 0.135 | 0.257 | 0.172 | 0.240 | 0.170 | 0.380 | 0.187 | 0.219 |
| Performant PLM Baselines |  |  |  |  |  |  |  |  |  |
| ProtBert Elnaggar et al. (2021) | ✗ | 0.188 | 0.279 | 0.464 | 0.456 | 0.234 | 0.408 | 0.859 | 0.838 |
| OntoProtein Zhang et al. (2022) | ✗ | 0.284 | 0.436 | 0.603 | 0.631 | 0.300 | 0.441 | 0.854 | 0.841 |
| Pretrained Backbone Baselines |  |  |  |  |  |  |  |  |  |
| ESM2-8M Rives et al. (2021) | ✓ | 0.154 | 0.284 | 0.410 | 0.394 | 0.187 | 0.373 | 0.477 | 0.468 |
| ESM2-35M Rives et al. (2021) | ✓ | 0.212 | 0.340 | 0.501 | 0.489 | 0.248 | 0.417 | 0.562 | 0.571 |
| Our models |  |  |  |  |  |  |  |  |  |
| ProtAC-PB | ✓ | 0.139 | 0.221 | 0.350 | 0.327 | 0.180 | 0.254 | 0.410 | 0.424 |
| ProtAC-ESM2-8M | ✓ | 0.239 | 0.354 | 0.454 | 0.423 | 0.307 | 0.431 | 0.579 | 0.558 |
| ProtAC-ESM2-35M | ✓ | 0.268 | 0.379 | 0.577 | 0.603 | 0.321 | 0.461 | 0.615 | 0.619 |

#### Stage Caption

We use 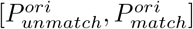 and 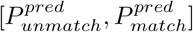 to represent the SAM Filter output of original and predicted sequence-annotation pair. The SAM Filter of our finetuned model determines whether the original annotation or the model-predicted annotation is closer to the corresponding protein sequence through two key conditions:

1. The model predicts that the compatibility between the annotation and this specific protein sequence has been predicted to be positive, indicating a successful match, *i*.*e*. 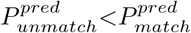.
2. The model-predicted annotation matches the protein sequence more closely than the original annotation, *i*.*e*. 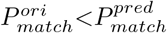.

Only when both conditions are satisfied will original annotation be replaced by model-predicted annotation. The cleaned dataset will replace previous noisy dataset in the next round of Stage Pretrain.

## 4 Experiments

### 4.1 Experimental Setups

#### Datasets and Annotations

We use **UniRef50** as of May 2018 for pre-training and captioning tasks, which contains 30.16 million protein sequences. **SwissProt**, updated to July 2023, contains 560,000 well-annotated sequences. We divide this database into two parts: 30,000 sequences are randomly selected for the test set, and the remaining approximately 530,000 sequences are used as the fine-tuning dataset. For the keyword prediction task, we further split the SwissProt test set in a 3:2 ratio, assigning 18,000 sequences to the training set and 12,000 sequences to the test set. The keywords associated with each sequence serve as labels to form the **Swiss-keyword** dataset. In the SwissProt caption task, we aggregate the newly updated sequences in SwissProt from 2023 to January 2024. We exclude sequences that overlap with sequences in the UniRef50 and SwissProt training datasets, resulting in a total of 458 sequences. This dataset is called the **Swiss-caption** dataset (see Appendix Tab.S3 for dataset details). We constructed annotation dictionaries of 7533 GO terms and 753 keywords, respectively (see Appendix B.1 for setup details).

#### Model and Training Configurations

We developed a ProteinBERT-based model and two ESM2-based models with different parameter versions: a small version using ESM2-8M and a basic version using ESM2-35M. Typically, we trained all models on eight A800 GPUs (time costs are shown in Appendix Fig.S6) with a training batch size of 256 for each model (equivalent to 32 proteins per GPU). We used the AdamW optimizer and an exponential learning rate scheduler, where the learning rate started at 1e-6, ramped up to 2e-5 during the first epoch, and then exponentially decreased back to 1e-6. Other settings are detailed in Appendix Tabs.S1 and S2.

#### Downstream Benchmark Tasks

To illustrate model capability in protein function annotation, we adopt two well-known benchmark tasks related to protein function, furthermore, we design a Keyword prediction task and a GO caption task using newly updated proteins in SwissProt:

- **Public Functional Annotation Tasks:** We adopted two established benchmarks introduced by DeepFRI Gligorijević et al. (2021), specifically for Enzyme Commission (EC) number prediction and Gene Ontology (GO) term prediction. The GO benchmarks are divided into three different branches: molecular function (abbreviated as GO-MF), biological process (GO-BP), and cellular component (GO-CC). Consistent with the task configuration in ProtST, we utilized dataset splitting to maintain a 95% sequence identity threshold for EC and GO predictions and applied a full model tuning strategy.
- **Keyword Prediction Task:** Keywords Magrane & Consortium (2011) are another important form of protein function annotation. To evaluate the transfer learning ability of the model, we designed a classification task. The sequence encoder and annotation decoder of the model pretrained on the GO prediction task were frozen. Only the application layer was finetuned for 200 epochs to evaluate the accuracy of the model in keyword prediction.
- **Gene Ontology Caption Task:** To evaluate the model’s ability to predict functions on never-seen sequences, we provide protein sequences and fully masked annotations from the Swiss-caption dataset as input to predict the corresponding GO annotations, which are then compared with newly curated GO terms from the SwissProt dataset.

### 4.2 Experimental Results

#### Our model shows SOTA performance in protein function prediction tasks

surpassing models with less than 100 million parameters and remaining competitive with larger PLMs. Tab.1 shows that our ProtAC-ESM2-35M ranks in the top three in four downstream tasks, achieves the highest score in GO-CC, and closely follows OntoProtein, a ProtBERT-based model with more than 400 million parameters, in GO-BP and GO-MF. In addition, our two ESM2-based models outperform their corresponding pretrained backbones in all tasks, and ProtAC-ESM2-8M not only outperforms ProtAC-PB, a model with similar number of parameters, but also surpasses ESM2-35M in three of the four tasks.

#### Models with pretrained weights exhibit superior performance compared to those trained from scratch

Fig.4a demonstrates that at an equivalent parameter count, the pretrained ProtAC-ESM2-8M model matches the SAM capability of the from-scratch ProtAC-PB model but significantly excels in GO prediction, indicating enhanced data cleaning efficiency through pretraining. Furthermore, as illustrated in Tab.2, comparisons between the 8M-cleaned and PB-cleaned versions reveal that, pretrained models consistently outperform their from-scratch counterparts.

**Figure 4:**
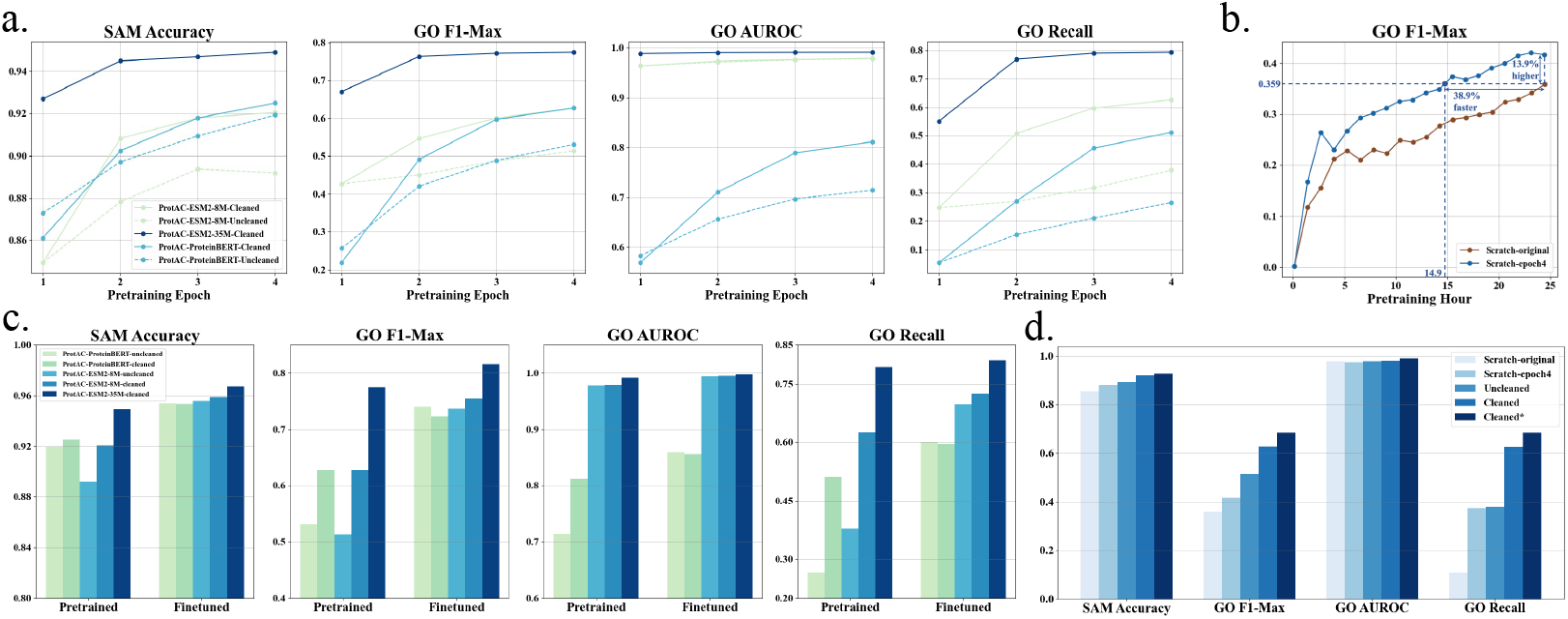
**(a)** Performance of different model versions during pretraining. The solid line represents the results of the model trained on the cleaned dataset, and the dashed line represents the results of the model trained on the original dataset. Each model was pretrained for four epochs and evaluated on the SwissProt test set after each epoch, using accuracy for SAM, F1-Max, AUROC, and recall for GO prediction. **(b)** Comparison of the improvement of pretraining on the original dataset and the cleaned dataset by GO prediction results. **(c)** Comparison between Finetuned and pretrained models. Cleaned: model trained on cleaned dataset; uncleaned: model trained on original dataset. The final results are presented after completing four rounds, where each round consists of pretraining the model for one epoch, followed by a finetuning phase of ten epochs. **(d)** Comparison of different pretraining strategies using ProtAC-ESM2-8M. Cleaned^*^: after the fourth epoch, the cleaned version is pretrained on the cleaned UniRef50 for one more full epoch.

#### Larger models exhibit superior performance

Fig.4a reveals that, among pretrained models, ProtAC-ESM2-35M outperforms ProtAC-ESM2-8M in both SAM and GO prediction, suggesting that increased model size enhances data cleaning efficacy. Additionally, Tab.2 shows that the 35M-cleaned model achieves the best outcomes across all four models, further supporting the conclusion that larger parameter sizes result in improved model performance.

#### Our curated dataset demonstrates efficacy

We applied Kaiming initialization He et al. (2015) to the 8M-version model, followed by one epoch of pretraining using both the original dataset (Scratch-original) and a dataset refined through four cleaning cycles (Scratch-epoch4). Validation is conducted on the SwissProt test set during training (See Fig.4b). Our findings indicate that the maximum *F*_*max*_ for Scratchepoch4 surpassed that of Scratch-original by 13.9%, and notably, Scratch-epoch4 achieved this benchmark *F*_*max*_ in 38.9% less training time. This evidence underscores the significant enhancement in the model’s protein function prediction capabilities attributable to our meticulously cleaned dataset.

**Table 2:**
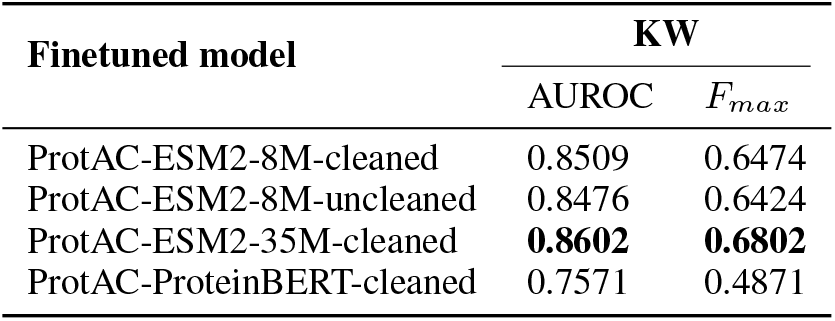
Keyword prediction results of ProtAC. In the main text, the four models listed in the table from top to bottom are referred to by the following abbreviated names: 8M-cleaned, 8M-uncleaned, 35M-cleaned, and PB-cleaned.

| Finetuned model | KW |  |
| --- | --- | --- |
| | AUROC | $F_{max}$ |
| ProtAC-ESM2-8M-cleaned | 0.8509 | 0.6474 |
| ProtAC-ESM2-8M-uncleaned | 0.8476 | 0.6424 |
| ProtAC-ESM2-35M-cleaned | **0.8602** | **0.6802** |
| ProtAC-ProteinBERT-cleaned | 0.7571 | 0.4871 |

#### Finetuned models surpass their pretrained counterparts in performance

Fig.4c illustrates that finetuned models outshine pretrained ones across all four metrics. For the 8M-version models, those finetuned on cleaned datasets exhibit superior performance to their uncleaned counterparts; conversely, for PB-version models, those finetuned on original datasets fare better. The results indicate substantial enhancements in SAM and GO prediction for models trained on original datasets following finetuning. Nevertheless, the performance gains from finetuning are more pronounced for models with fewer parameters than for those with a larger parameter count.

#### Our cleaning strategy yields positive results

Fig.4a demonstrates that pretraining models on cleaned datasets enhances SAM and GO prediction capabilities compared to models pretrained on noisy datasets, indicating the effectiveness of our data cleaning approach for protein function prediction. Tab.2 further supports this observation by showing that models trained on cleaned datasets outperform those trained on original datasets of the same architecture. Additionally, our training approach utilizing different datasets is highlighted in Fig.4d, where Scratch-epoch4 outperforms Scratch-original in three out of four metrics, showcasing the continuous improvement in dataset quality facilitated by our cleaning strategy. Notably, the performance of the Cleaned^*^ model excels in all four metrics, indicating that extended pretraining enhances the outcomes of our cleaning strategy (more comparison is shown in Appendix B.2).

#### Data caption results show significant improvement

Fig.5 shows that after training, our model’s prediction performance in Swiss-caption has been significantly improved, and is far better than the original functional annotations of the dataset. Finetuned models generally outshine pretrained models in GO caption performance. Larger parameter sizes contribute to better performance, pretrained models outperform scratch-trained models, and the ProtAC-ESM2-35M-finetuned model stands out as the top performer among all versions. The quantities of sequences in UniRef50 cleaned by each model in every round are also delineated in Appendix Fig.S5.

**Figure 5:**
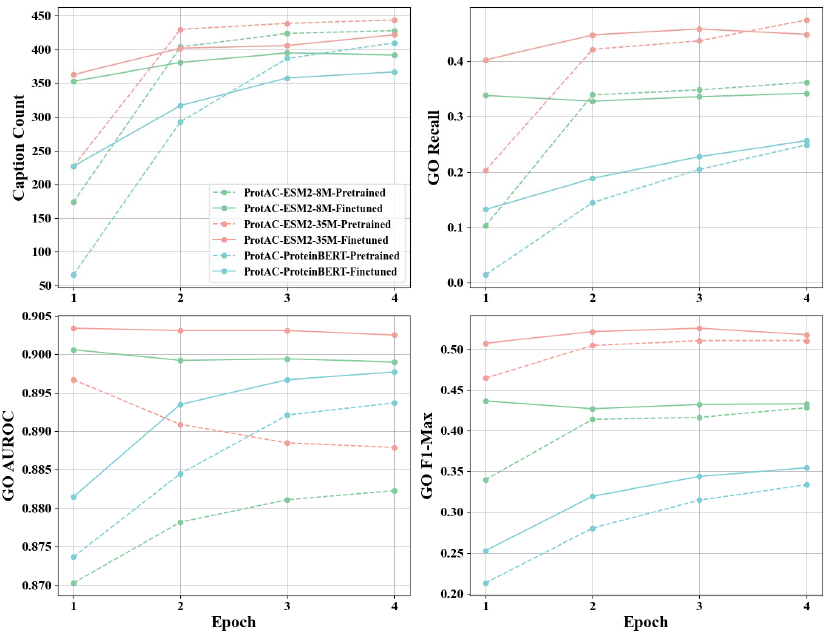
Protein function caption results across three model versions. The function of 418 sequences out of 458 has been captioned in newly updated SwissProt dataset. Notably, original GO yields an F1-Max of 0.0695, an AUROC of 0.1099, and a recall of 0.0344.

### 4.3 Biological Analysis

#### Manual evaluation (GO annotation comparison for same protein)

To further validate the biological significance of the data cleaning results for ProtAC, we conduct a manual verification of the GO annotations before and after cleaning on a sampled set of protein data. The verification process involves the following steps: (1) random extracting 30 UniRef clusters with newly added GO terms after cleaning; (2) querying UniProt (as of 2024-07-30) for the existing GO annotations and associated family and domain information for sequences in these clusters; (3) verifying whether the new GO terms added by ProtAC correspond to updates in existing databases or to their ancestor or child terms, and assessing whether the family and domain information provides supportive evidence for the newly added GO terms. Among the randomly sampled 30 clusters, those that are either deprecated or have an excess of member sequences that can not be manually verified are excluded from the analysis. The newly added GO terms for the selected five clusters (Appendix Tab.S5; Fig. 6a) are supported by evidence. For example, UniRef50 A0A1I4VGP3 (Fig. 6a, top) contains a member sequence, A0A1I4VGP3, which originally lacked corresponding GO annotations. After data cleaning with ProtAC, GO:0005886 is added in the first three rounds, and GO:0055085 is added in the fourth round. Upon review, the current GO annotation for this cluster in UniProt is GO:0016020. Both GO:0005886 and GO:0055085 are child terms of GO:0016020. Further investigation into the family and domain information for A0A1I4VGP3 reveals that it matches the IPR002549 family in the InterPro database. This family, known as the Transmembrane protein TqsA-like family, regulates quorum-sensing signal transmission by either enhancing the secretion of autoinducer-2 (AI-2) or inhibiting its uptake. This information suggests the potential inclusion of GO:0005886 and GO:0055085, indicating that ProtAC may enhance the granularity of protein annotations. Similarly, UniRef50 A0A1I1LTJ9 (Fig. 6a, bottom) contains two member sequences, both of which are annotated with GO:0016020 in the latest version of UniProt. Following ProtAC curation, GO:0016020 is added in the first round; in the second and third rounds, GO:0009881 and GO:0006355 are added; in the fourth round, GO:0006355 is removed, and GO:0030435 and GO:0046872 are added. Upon analysis, GO:0009881 and GO:0046872 are identified as co-occurring terms with GO:0016020, and these annotations are also supported by evidence from family and domain databases for the two member sequences of UniRef50 A0A1I1LTJ9.

**Figure 6:**
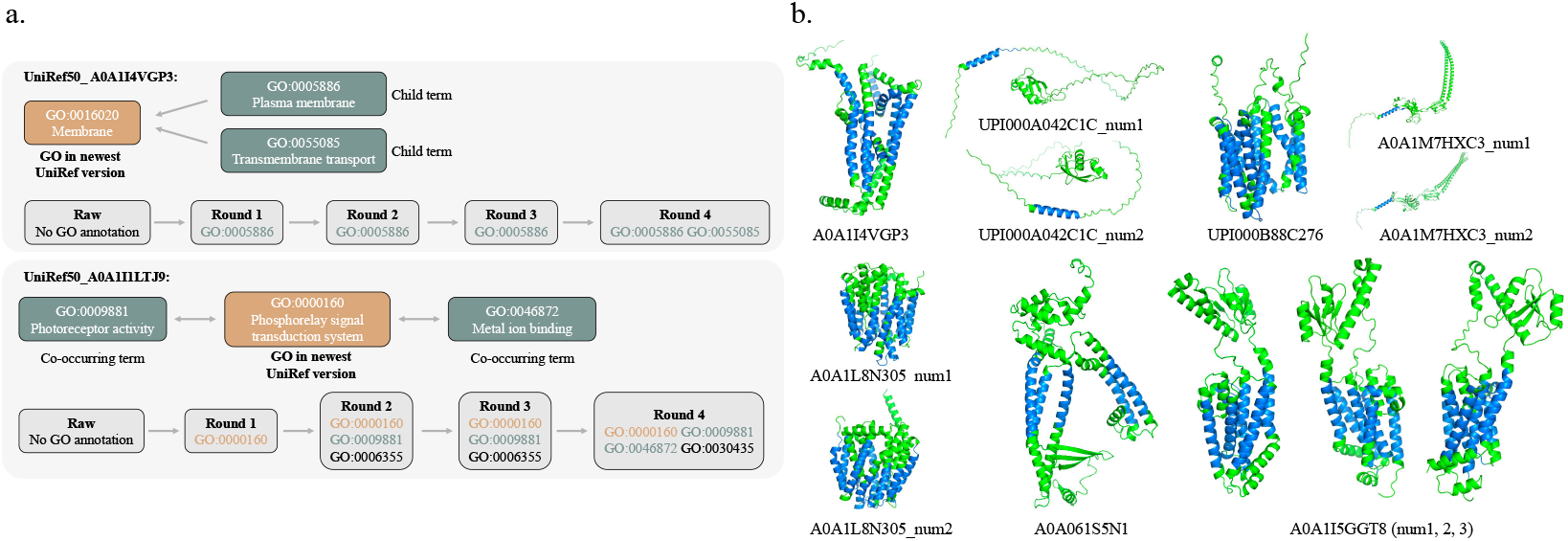
**(a)** Examples of GO variation analysis for the same protein in each cleaning round. We use two color to denote the GO annotations **in the newest UniRef** and **supported by family databases**. The child term would be a more specific term; the co-occurring terms would be coannotated for the same protein or gene. **(b)** Predicted structures of transmembrane proteins. We use two different colors to distinguish between **transmembrane domains** and **other sequences**.

#### Manual evaluation (Protein comparison for same GO)

After conducting a manual verification of the same protein’s GO annotations before and after cleaning with ProtAC, we further analyze whether the protein sequences in clusters annotated with “transmembrane” related terms contained transmembrane regions. We first filter 152 GO annotations containing the term “transmembrane” from a total of 7,533 GO annotations in the GO dictionary. Subsequently, we randomly sample 20 clusters that contain neither of these 152 GO annotations before using ProtAC and contain some of these 152 GO annotations after the cleaning process. We use the Phobius Protein Functional Analysis tool (https://www.ebi.ac.uk/jdispatcher/pfa/phobius) to predict transmembrane regions in all member sequences of the selected clusters. Among the 20 sampled clusters, 11 clusters are no longer present in UniRef50, and one cluster’s sequences do not predict any transmembrane regions (Fig.6b). Transmembrane regions are predicted in the remaining 8 clusters. These results further underscore the biological relevance of ProtAC’s performance in refining GO annotations (more biological analysis shown in Appendix B.3).

### 4.4 Ablation Study

In the Experimental Results section, we have conducted comprehensive comparisons regarding the impact of the cleaned dataset, various cleaning strategies, and different model sizes on training effectiveness. Hence, in this section, we primarily investigate the influence of three losses on model performance. We pretrain ProtAC-PB for one epoch.

Tab.3 shows that, eliminating ℒ _*SAM*_ and ℒ _*MLM*_, which are not directly related to function prediction, enhances the *F*_*max*_ of pretraining GO and downstream Keyword prediction. However, removing any of the three losses significantly reduces the *F*_*max*_ for GO captioning and downstream EC tasks. This underscores the critical importance of synthetic data filtration for model training Shumailov et al. (2024).

**Table 3:** Ablation study on three losses using pretrained models. We show the *F*_*max*_ for four function prediction tasks. *Abbr*., **Pre. GO**: GO prediction task in pretraining; **Cap. GO**: GO caption task. Notably, Pre. GO is dependent on ℒ _*AP*_ for its functionality.

| Model | Pre. GO | Cap. GO | EC | KW |
| --- | --- | --- | --- | --- |
| Full loss | 0.2179 | 0.2132 | 0.4026 | 0.4282 |
| w/o $\mathcal{L}_{SAM}$ | 0.2488 | 0.0506 | 0.3348 | 0.4311 |
| w/o $\mathcal{L}_{MLM}$ | 0.3974 | 0.1114 | 0.3857 | 0.4607 |
| w/o $\mathcal{L}_{AP}$ | / | 0.0344 | 0.3477 | 0.3785 |

## 5 Conclusion

We introduce ProtAC, a recursive cleaning framework that uses protein multimodal learning to optimize noisy annotations in large-scale protein datasets while enhancing the learning capabilities of PLMs. We develop a model with fewer than 100M parameters that achieves SOTA results on multiple function-related downstream tasks while also cleaning up a high-quality protein dataset.

However, limited by computational resources, our learning framework still has significant room for improvement. For instance, a protein’s structure dictates its functionality, making structural information crucial for models to learn functional annotations accurately. Moreover, the vast research literature related to protein functions contains an abundance of extractable feature. Therefore, incorporating other modalities of information into our research is one of our future goals. In addition, this work has already demonstrated the immense potential of larger-scale PLMs, and we plan to explore the boundaries of protein function research further by introducing higher parameter-level PLMs in future studies. We also anticipate that the achievements of this work, including the curated dataset, can be applied to other tasks in protein representation learning, *e*.*g*. large-scale pretraining of protein models or *de novo* protein design.

## A More Related Work

### A.1 Large-scale Protein Datasets

Large datasets can help to improve the scalability of the model and provide a more comprehensive representation of the underlying data distribution. Their comprehensive coverage of protein sequences from thousands of species enables the development of more robust and generalizable computational models. UniProt dataset Magrane & Consortium (2011) offers unparalleled representation of protein diversity sith over 200 million sequences from more than 20,000 species. This allows researchers to draw insights from a huge array of proteins. In contrast to specialized databases like Protein Data Bank(PDB) Burley et al. (2019) and Gene Expression Omnibus (GEO) Clough & Barrett (2016) which focus on narrow data types, UniProt consolidates information from genomic, proteomic, and functional sources. This multi-modal view facilitates analysis of proteins from numerous angles. UniRef90, a protein sequence database that clusters sequences at 90 percents identity Suzek et al. (2007), further enhances UniProt by reducing redundancy through sequence clustering. Similarly, UniRef50 is built by clustering UniRef90 seed sequences that have at least 50% sequence identity to and 80% overlap with the longest sequence in the cluster. The non-redundant sequences improve annotation quality and search efficiency. Regular updates also ensure researchers have access to the latest discoveries. By leveraging the scale and diversity of data in UniProt and UniRef, scientists can gain a deeper understanding of proteins and their many functions. These large-scale databases are foundational to modern bioinformatics.

### A.2 Protein Annotation Description

Comprehensively describing the diverse functions of proteins is critical for interpreting their roles in biological systems. While several annotation types exist, GO terms Ashburner et al. (2000) and Keywords Magrane & Consortium (2011) are especially valuable. GO terms from the Gene Ontology allow consistent representation of molecular functions, biological processes, and cellular components across species. Their widespread use enables both granular annotation of individual proteins and higher-level pathway enrichment analysis. This dual utility makes GO terms a fundamental tool for functional genomics research Huang et al. (2009). Keywords from UniProtKB similarly provide standardized vocabulary for protein functions. Manually curated for Swiss-Prot and automatically assigned for TrEMBL, Keywords capture multifaceted functional aspects in a structured ontology Magrane & Consortium (2011). The hierarchical organization into categories like molecular function and biological process aids literature indexing and database searching. By consolidating expert knowledge into controlled terminologies, GO terms and Keywords empower accurate computational analysis and biological interpretation. Their adoption throughout public bioinformatics databases highlights the indispensable role protein function annotation plays in translating sequence data into actionable knowledge.

## B Additional Experimental Results

### B.1 Functional Annotation Setups

To construct an exhaustive Gene Ontology (GO) dictionary, we enumerated the number of occurrences of all GO terms in the UniRef and SwissProt datasets. We considered only those GO terms that appeared 100 times or more, resulting in a dictionary of 7533 terms (see Tab.S4 for composition analysis of the GO dictionary). The keyword dictionary included all keywords that appeared in the Swiss-keyword dataset, totaling 753 keywords. See Appendix A.2 for an overview of GO and keywords.

### B.2 Additional Experiments Related TO Methodology

#### B.2.1 From-scratch Training Setups

We developed two distinct parametric models based on ProteinBERT, labeled as “ProtAC-PB-small” and “ProtAC-PB-base”. The “small” variant incorporates 6 layers and 4 attention heads, while the “base” model comprises 12 layers and 8 attention heads. These models underwent training on eight A800 GPUs, utilizing an AdamW optimizer in conjunction with a learning rate scheduler that includes a warm-up step followed by exponential decay. The initial learning rate was set to 1e-6, which was increased to 3e-4 during the warm-up phase, before being exponentially decreased back to 1e-6. The decay factor for the learning rate was maintained at 0.9, with the warm-up period lasting for 1 epoch. Moreover, the models were trained with a batch size of 128.

#### B.2.2 Cleaning Strategy Works FOR Model Trained From-scratch

Fig. S1(a) illustrates the evolution of the base model’s annotation prediction F1-score throughout the pretraining stage over three rounds. The graph demonstrates a progressive increase in the growth rate of the model’s F1-score curve through successive cleanup cycles, coupled with a significant improvement in the peak value achieved. This pattern underscores the effectiveness of our datacleaning strategy in enhancing the model’s learning performance.

#### B.2.3 Continuous Cleaning Strategy

We explored two distinct data cleaning methodologies: the continue caption strategy and the notcontinue caption strategy, obtaining valuable insights from both. Our validation approach comprised several steps: Initially, in Epoch 1, the small model was pretrained and fine-tuned using the uniref90 (original) dataset, followed by data cleaning to produce uniref90 (epoch1). Subsequently, in Epoch 2, the model was pretrained and fine-tuned on uniref90 (epoch1), and data cleaning was performed against both uniref90 (original) and uniref90 (epoch1) to create uniref90 (epoch2-nocontinue) and uniref90 (epoch2-continue). In Epoch 3, the model underwent pretraining and its training metrics were evaluated on uniref90 (epoch2-continue) and uniref90 (epoch2-nocontinue), respectively. Fig. S1(b) delineates the comparative analysis of the annotation prediction performance of the small model utilizing the no-continue caption and continue caption strategies. This figure illustrates two distinct curves that trace the maximum F1-score trajectories of the small model in Epoch 3, following pretraining on the uniref90 (epoch2-continue) and uniref90 (epoch2-nocontinue) datasets, correspondingly. The data clearly indicates that pretraining on the uniref90 (epoch2-continue) dataset results in a higher F1-score, thus underscoring the superior training effectiveness of the continue caption strategy. Based on these findings, the continue caption strategy was consistently employed for both training and data cleaning throughout our investigation.

#### B.2.4 Impact OF Model Parameters ON Cleaning Process Time AND Data Cleaning Efficacy

Fig. S1(c) provides a comparative analysis of the temporal investment and testing performance across four iterative cleaning cycles for the small model and three cycles for the base model. The aggregate time spent per epoch by both models was meticulously recorded, and the refined models’ annotation prediction proficiency was assessed using the SwissProt test set, employing the maximum F1-score metric. The outcomes of this assessment are depicted in a line graph. The analysis indicates that the duration required for the small model to undergo four cleaning cycles is approximately two-thirds that of the base model’s completion of three cycles. Furthermore, the small model exhibits superior F1-score and AUC values relative to the base model, suggesting that the small model achieves improved learning efficiency and outcomes over multiple iterations within a reduced timeframe.

### B.3 More Biological Analysis

#### B.3.1 Comparison OF Visualization Results: Original VS. Cleaned Dataset Model Training

To verify that the cleaned dataset improves the performance of our model, we compared the clustering results of the model trained by the original Uniref vs. the model trained by the cleaned Uniref. We select protein sequences in *cellular component* GO domain in the Swissprot-test dataset. We then apply t-SNE to visualize the clustering of the seq embeddings from the model’s sequence encoder. The clustering results (depicted in Fig. S2) demonstrate that embeddings derived from the model trained on the cleaned dataset exhibit significantly better coherence and separation. Taking three distinct subcellular compartments, namely cytosol (GO:0005829), extracellular region (GO:0005576), and nucleus (GO:0005634), which are spatially separated, as an example. We visualized the protein data that only contains annotations for one of these three compartments. It is evident that the embeddings obtained from models trained on the original data exhibit significant overlap, whereas the embeddings obtained from models trained on the cleaned data are distinctly separated from each other. This indicates that the cleaned dataset can enhance the model representative ability.

#### B.3.2 Comparative Visualization OF Original AND Cleaned Sequence EMBEDDINGS

We conduct visualization analyses on the original dataset and the cleaned dataset, verifying the improvement of data quality. We first extract the sequences where the GO terms are revised after cleaning and then select a list of GO terms with a high number of occurrences. To ensure a fair comparison, we choose the model trained on SwissProt for embedding extraction. Then, we apply t-SNE to obtain the clustering outcomes of the original sequence embeddings vs. cleaned sequence embeddings. Fig. S3 reveals that the embeddings from the cleaned dataset result in markedly improved clustering, characterized by enhanced grouping and distinctiveness for sequences associated with the same GO terms. This outcome demonstrates the improved quality of the cleaned dataset.

#### B.3.3 Comparision OF THE Biophysical Embeddings OF Amino Acids

The biophysical properties of amino acids, e.g., hydrophobicity, aromaticity, and charge, are widely recognized to profoundly impact the structural configurations of proteins. We visualize the biophysical properties of amino acids. Fig. S4 illustrates the comparative analysis of clustering outcomes between the model trained on the original dataset and one trained on the cleaned dataset. The findings indicate that models trained on the cleaned dataset show a slight improvement in clustering performance. In this regard, the distances between similar amino acids are more compact compared to before the cleaning process (e.g., hydrophobic (aromatic), polar neutral, positive amino acids), while there is a clear separation on the plane between hydrophobic amino acids and hydrophilic amino acids (positive and negative amino acids). This suggests that data cleaning significantly contributes to the model’s ability to categorize the underlying biophysical characteristics of amino acids more effectively.

**Figure S1:**
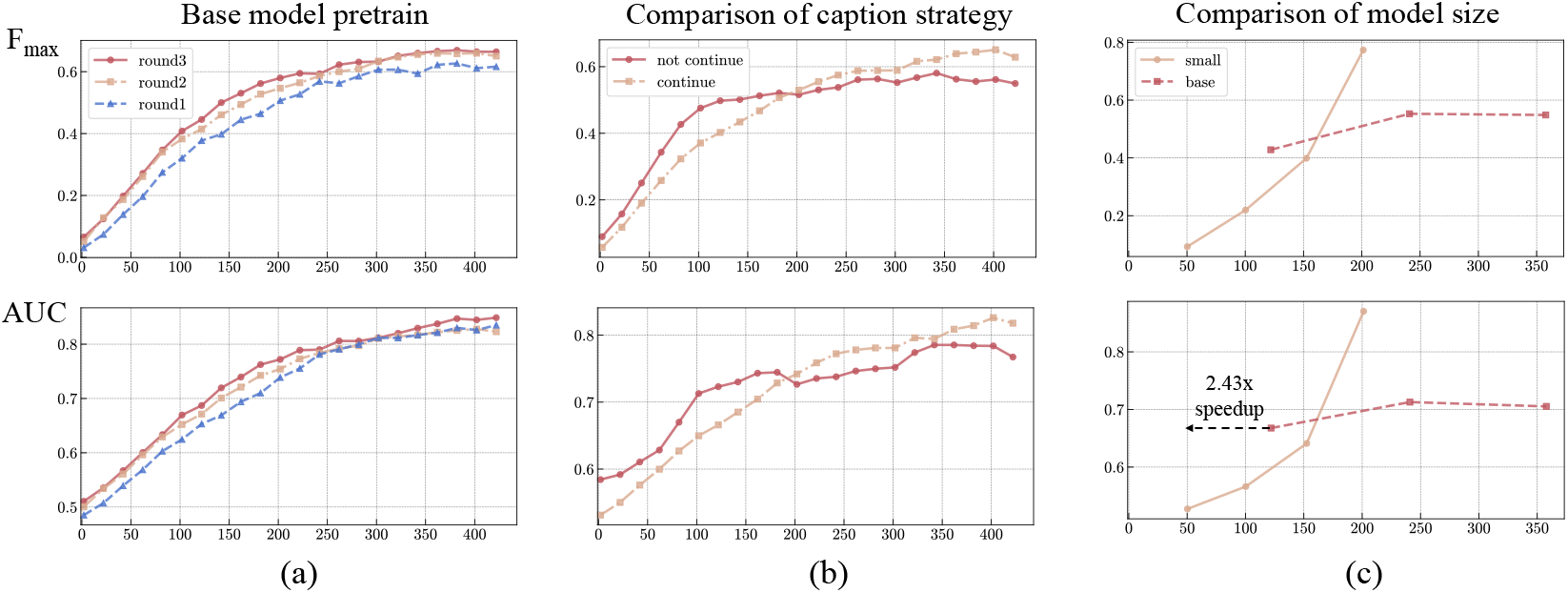
Visualisation of training results. The horizontal axis measures the training steps, where each step encompasses 1600 batches, and the vertical axis denotes the maximum F1-score achieved by the model in annotation prediction. (a) Annotation prediction curves of the base-model in the training phase of round1-3. (b) Comparison of the effects of different caption strategies on model training results. (c) Comparison of the learning effects of models at different scales.

**Figure S2:**
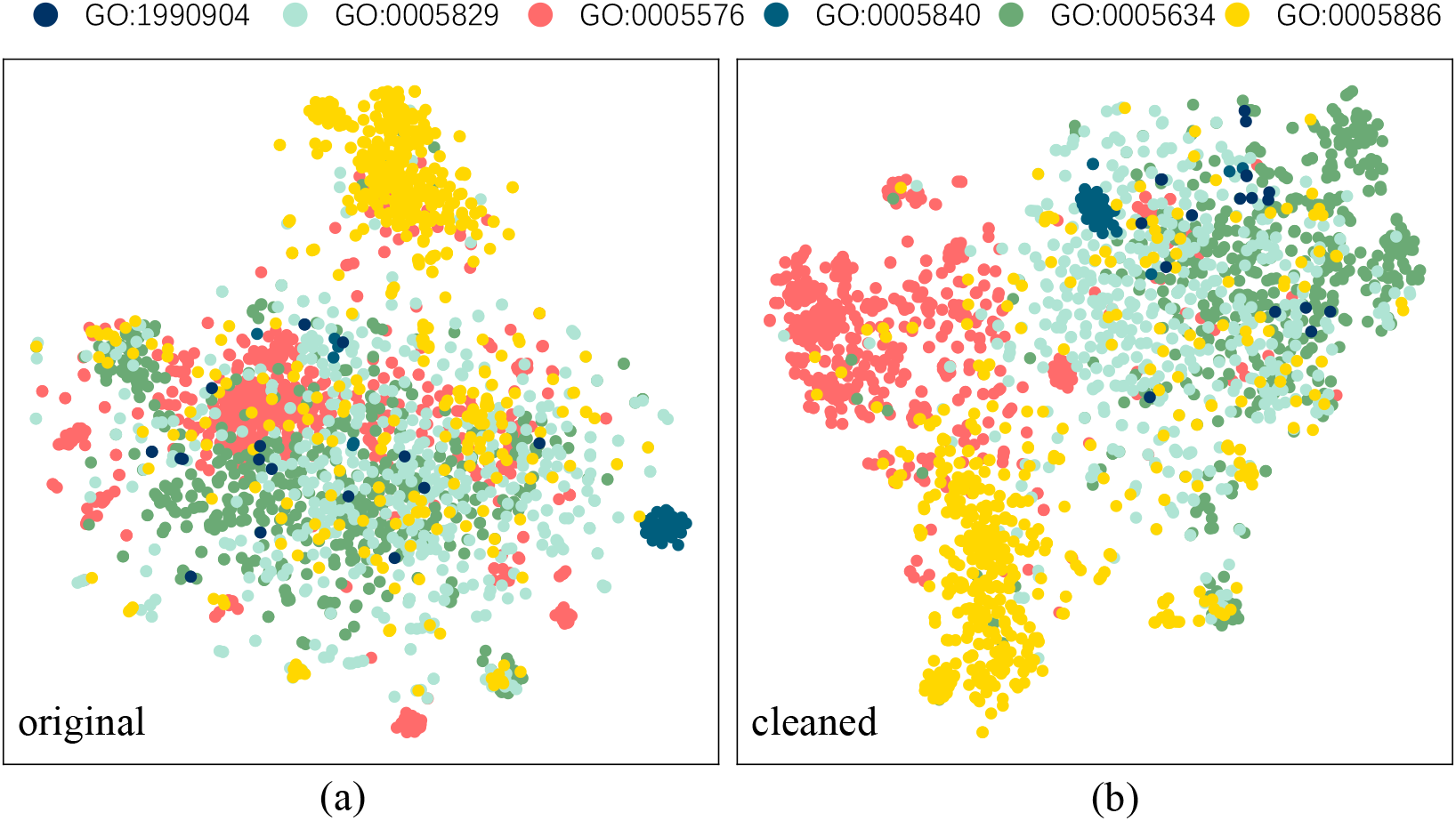
Visualization results by using the model trained on original (a) vs. cleaned dataset (b). The improved clustering outcome demonstrates that the cleaned dataset enhances the model representative ability.

**Table S1:** ProtAC model details.

| model version | seq encoder | anno encoder | layer | head | parameters |
| --- | --- | --- | --- | --- | --- |
| ProtAC-ProteinBERT | ProteinBERT local part | ProteinBERT global part | 6 | 4 | 27M |
| ProtAC-ESM2-8M | ESM-8M |  | 6 | 4 | 29M |
| ProtAC-ESM2-35M | ESM-35M |  | 12 | 8 | 78.56M |

**Figure S3:**
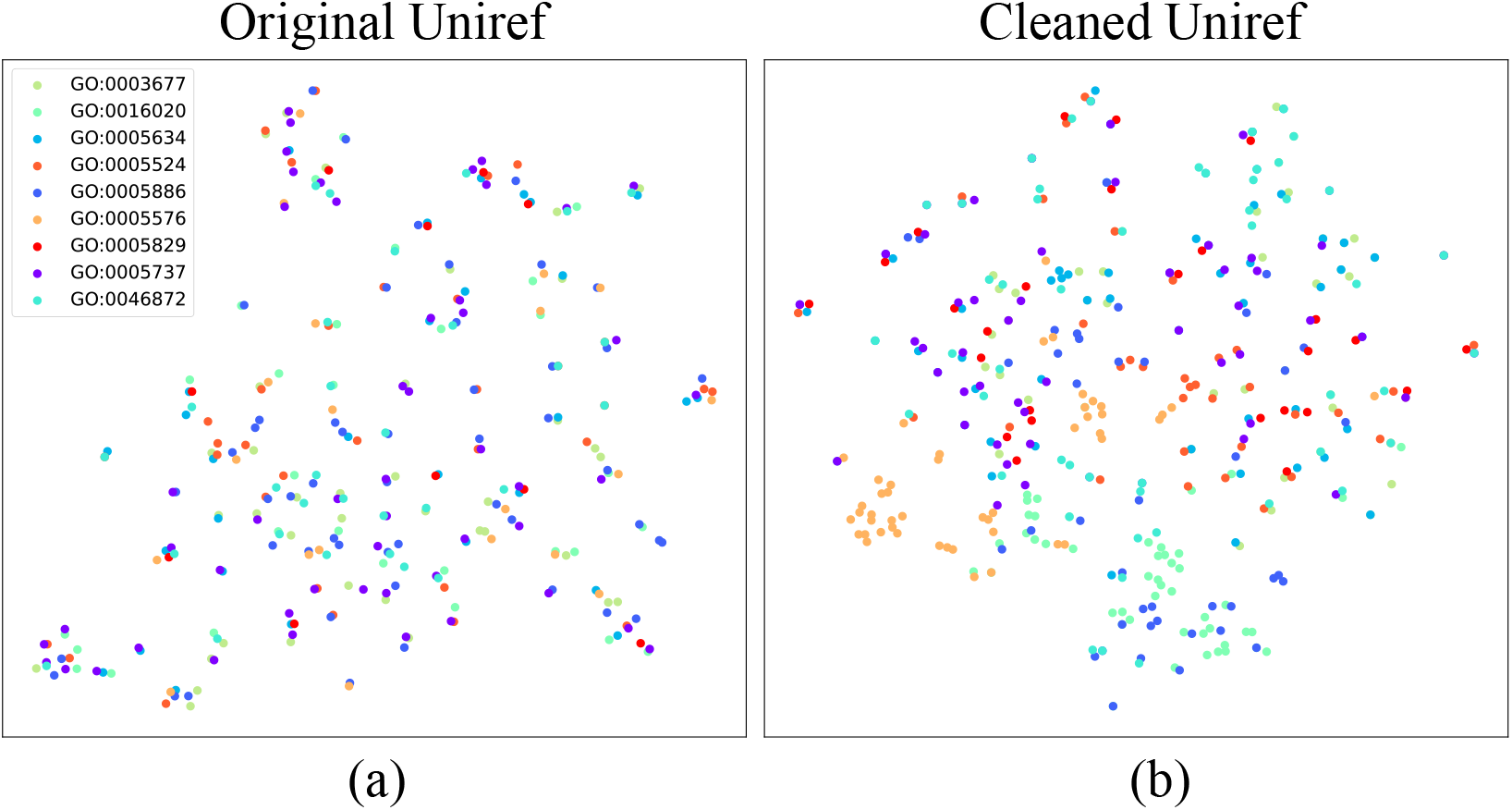
Visualization results of original vs. cleaned sequence embeddings. The cleaned dataset achieves better clustering results which validates the improved quality of the cleaned data.

**Figure S4:**
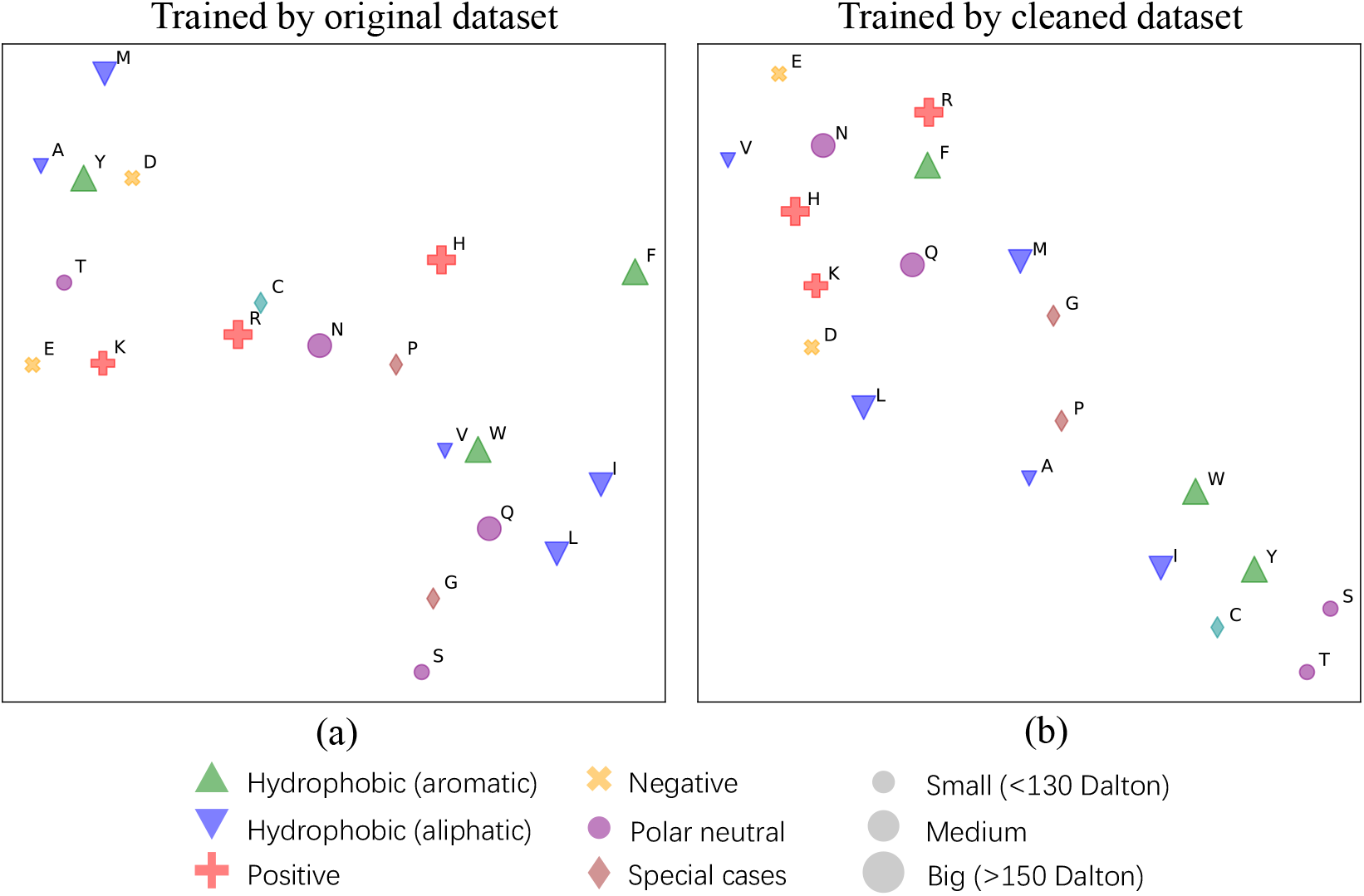
Biophysical embedding of amino acids.

**Figure S5:**
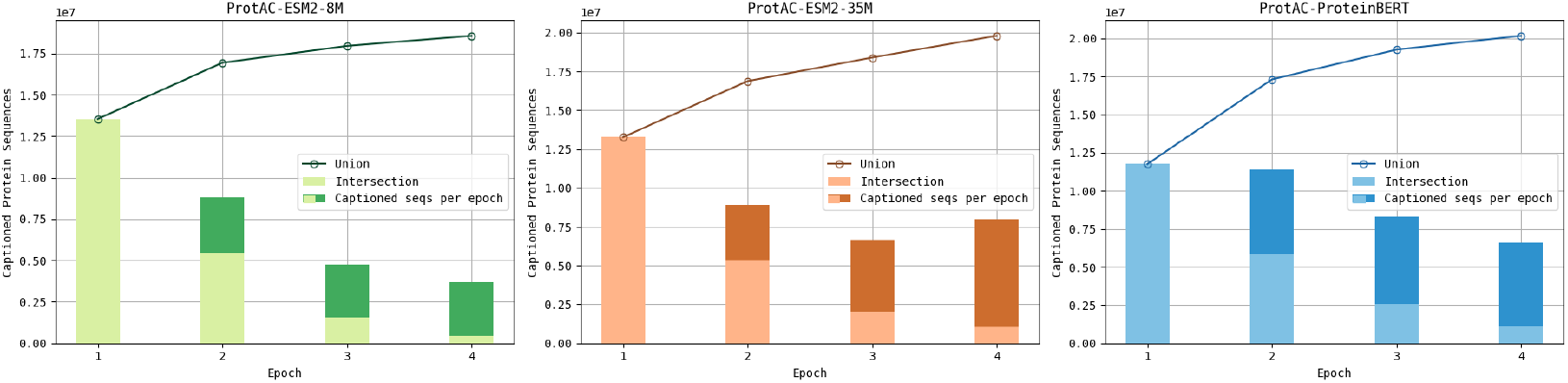
Captioned sequences count in UniRef50

**Figure S6:**
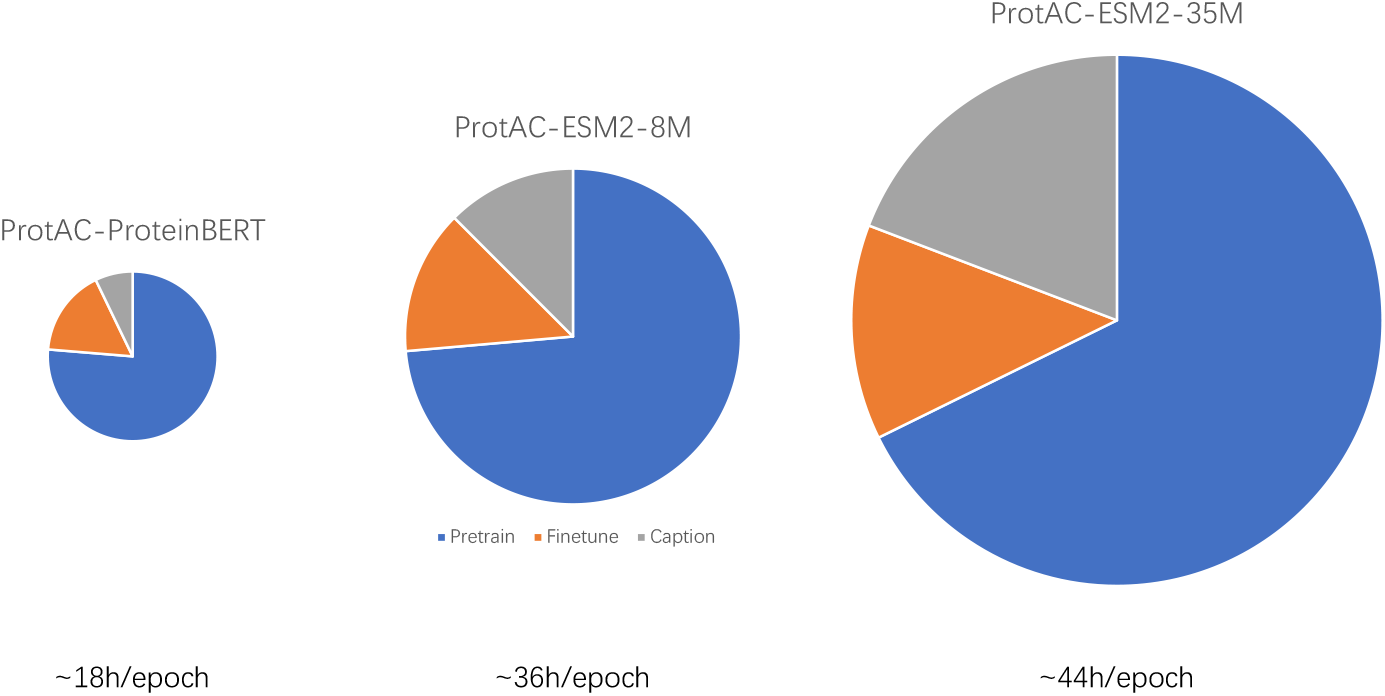
Time cost of each model version for one cleaning epoch/round

**Table S2:** ProtAC training details.

| Hyperparameter | Value |
| --- | --- |
| Input sequence length | 512 |
| Optimizer | AdamW |
| Scheduler | ExponentialLR |
| Pretraining epoch per round | 0.2 (ESM-8M)<br>1 (ProteinBERT&ESM-35M) |
| Finetuning epoch per round | 10 |
| Round | 4 |
| Pretraining dataset | UniRef50_2018_5 |
| Finetuning dataset | SwissProt_2023 |
| Init LR | 2e-5 |
| LR decay rate | 0.9 |
| Min LR | 1e-6 |
| Training batch size (8GPUs) | 512 |

**Table S3:** Datasets details.

| UniRef50_2018 | SwissProt |  | SwissProt(keyword) |  | SwissProt(caption) |
| --- | --- | --- | --- | --- | --- |
|  | trainset | testset | trainset | testset |  |
| ~30.16M | ~530K | ~30K | 18K | 12K | 458 |

**Table S4:** GO dictionary details.

| GO Classification | Amount |
| --- | --- |
| CC (Cellular Component) | 962 |
| BP (Biological Process) | 3346 |
| MF (Molecular Function) | 3225 |
| Total | 7533 |

**Table S5:**
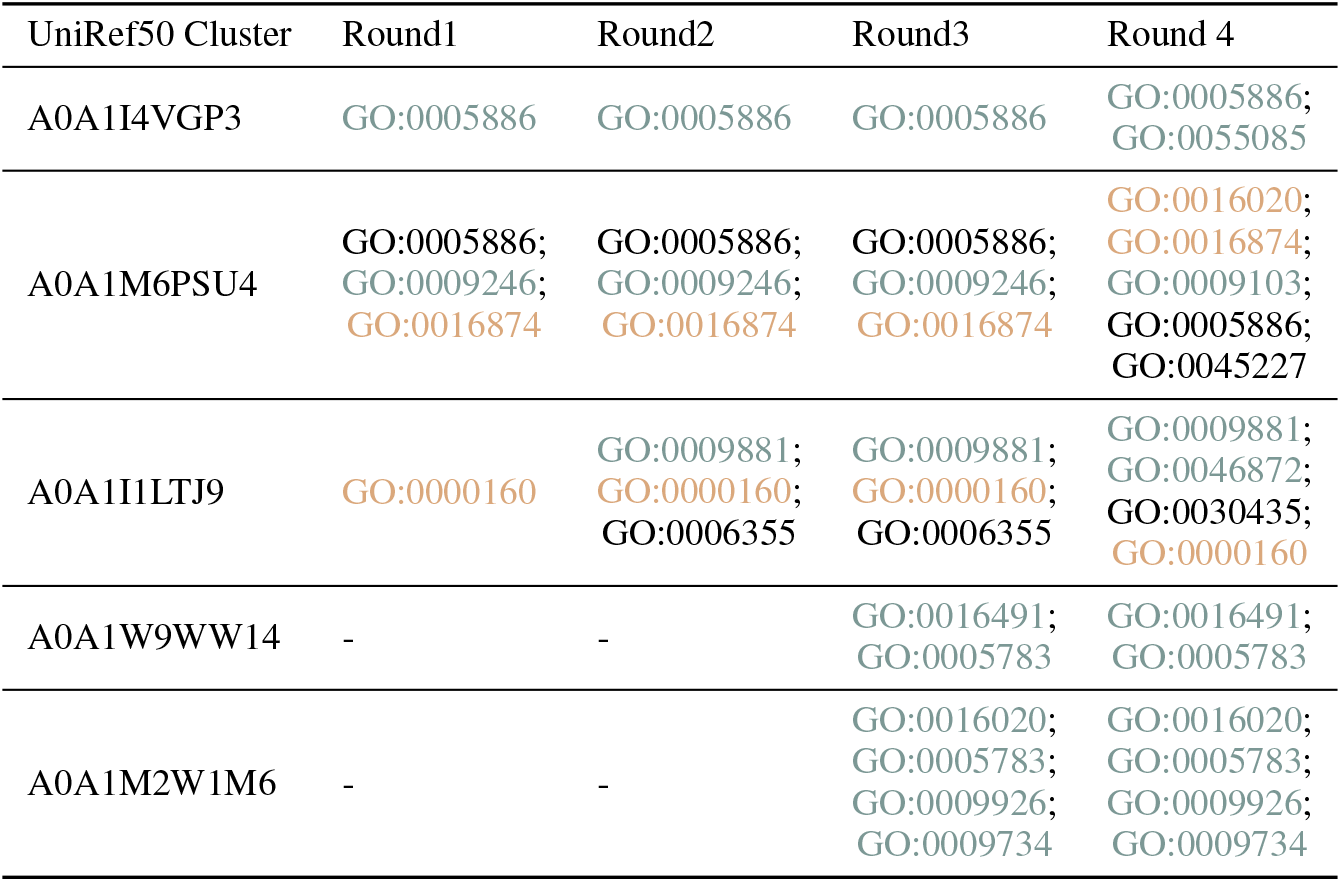
GO comparison for same proteins. We use two color to denote the GO annotations **in the newest UniRef** and **supported by family databases**.

## Notes

### Competing Interest Statement

The authors have declared no competing interest.

## References

Josh Abramson, Jonas Adler, Jack Dunger, Richard Evans, Tim Green, Alexander Pritzel, Olaf Ronneberger, Lindsay Willmore, Andrew J Ballard, Joshua Bambrick, et al. Accurate structure prediction of biomolecular interactions with alphafold 3. Nature, pp. 1–3, 2024.

Michael Ashburner, Catherine A Ball, Judith A Blake, David Botstein, Heather Butler, J Michael Cherry, Allan P Davis, Kara Dolinski, Selina S Dwight, Janan T Eppig, et al. Gene ontology: tool for the unification of biology. Nature genetics, 25(1):25–29, 2000.

Minkyung Baek, Frank DiMaio, Ivan Anishchenko, Justas Dauparas, Sergey Ovchinnikov, Gyu Rie Lee, Jue Wang, Qian Cong, Lisa N Kinch, R Dustin Schaeffer, et al. Accurate prediction of protein structures and interactions using a three-track neural network. Science, 373(6557):871– 876, 2021.

Emmanuel Boutet, Damien Lieberherr, Michael Tognolli, Michel Schneider, and Amos Bairoch. Uniprotkb/swiss-prot: the manually annotated section of the uniprot knowledgebase. In Plant bioinformatics: methods and protocols, pp. 89–112. Springer, 2007.

Nadav Brandes, Dan Ofer, Yam Peleg, Nadav Rappoport, and Michal Linial. Proteinbert: a universal deep-learning model of protein sequence and function. Bioinformatics, 38(8):2102–2110, 2022.

Stephen K Burley, Helen M Berman, Gerard J Kleywegt, John L Markley, Haruki Nakamura, and Sameer Velankar. Protein data bank (pdb): the single global macromolecular structure archive. Protein crystallography: methods and protocols, pp. 627–641, 2017.

Stephen K Burley, Helen M Berman, Charmi Bhikadiya, Chunxiao Bi, Li Chen, Luigi Di Costanzo, Cole Christie, Ken Dalenberg, Jose M Duarte, Shuchismita Dutta, et al. Rcsb protein data bank: biological macromolecular structures enabling research and education in fundamental biology, biomedicine, biotechnology and energy. Nucleic acids research, 47(D1):D464–D474, 2019.

Bo Chen, Xingyi Cheng, Pan Li, Yangli-ao Geng, Jing Gong, Shen Li, Zhilei Bei, Xu Tan, Boyan Wang, Xin Zeng, et al. xtrimopglm: unified 100b-scale pre-trained transformer for deciphering the language of protein. arXiv preprint arXiv:2401.06199, 2024.

Emily Clough and Tanya Barrett. The gene expression omnibus database. Statistical Genomics: Methods and Protocols, pp. 93–110, 2016.

UniProt Consortium. Uniprot: a worldwide hub of protein knowledge. Nucleic acids research, 47 (D1):D506–D515, 2019.

Jacob Devlin, Ming-Wei Chang, Kenton Lee, and Kristina Toutanova. Bert: Pre-training of deep bidirectional transformers for language understanding. arXiv preprint 1810.04805, 2018.

Ahmed Elnaggar, Michael Heinzinger, Christian Dallago, Ghalia Rehawi, Yu Wang, Llion Jones, Tom Gibbs, Tamas Feher, Christoph Angerer, Martin Steinegger, et al. Prottrans: Toward understanding the language of life through self-supervised learning. IEEE transactions on pattern analysis and machine intelligence, 44(10):7112–7127, 2021.

Noelia Ferruz, Steffen Schmidt, and Birte Höcker. Protgpt2 is a deep unsupervised language model for protein design. Nature Communications, Jul 2022. doi: 10.1038/s41467-022-32007-7. URL 10.1038/s41467-022-32007-7.

Vladimir Gligorijević, P Douglas Renfrew, Tomasz Kosciolek, Julia Koehler Leman, Daniel Berenberg, Tommi Vatanen, Chris Chandler, Bryn C Taylor, Ian M Fisk, Hera Vlamakis, et al. Structurebased protein function prediction using graph convolutional networks. Nature communications, 12(1):3168, 2021.

Kaiming He, Xiangyu Zhang, Shaoqing Ren, and Jian Sun. Delving deep into rectifiers: Surpassing human-level performance on imagenet classification. In Proceedings of the IEEE international conference on computer vision, pp. 1026–1034, 2015.

Geoffrey Hinton, Oriol Vinyals, and Jeff Dean. Distilling the knowledge in a neural network. arXiv preprint arXiv:1503.02531, 2015.

Da Wei Huang, Brad T Sherman, and Richard A Lempicki. Systematic and integrative analysis of large gene lists using david bioinformatics resources. Nature protocols, 4(1):44–57, 2009.

John Jumper, Richard Evans, Alexander Pritzel, Tim Green, Michael Figurnov, Olaf Ronneberger, Kathryn Tunyasuvunakool, Russ Bates, Augustin Žídek, Anna Potapenko, et al. Highly accurate protein structure prediction with alphafold. nature, 596(7873):583–589, 2021.

Maxat Kulmanov, Mohammed Asif Khan, and Robert Hoehndorf. Deepgo: predicting protein functions from sequence and interactions using a deep ontology-aware classifier. Bioinformatics, 34 (4):660–668, 2018.

Junnan Li, Ramprasaath Selvaraju, Akhilesh Gotmare, Shafiq Joty, Caiming Xiong, and Steven Chu Hong Hoi. Align before fuse: Vision and language representation learning with momentum distillation. Advances in neural information processing systems, 34:9694–9705, 2021.

Junnan Li, Dongxu Li, Caiming Xiong, and Steven Hoi. Blip: Bootstrapping language-image pretraining for unified vision-language understanding and generation. In International conference on machine learning, pp. 12888–12900. PMLR, 2022.

Tsung-Yi Lin, Priya Goyal, Ross Girshick, Kaiming He, and Piotr Dollár. Focal loss for dense object detection. In Proceedings of the IEEE international conference on computer vision, pp. 2980–2988, 2017.

Zeming Lin, Halil Akin, Roshan Rao, Brian Hie, Zhongkai Zhu, Wenting Lu, Nikita Smetanin, Robert Verkuil, Ori Kabeli, Yaniv Shmueli, et al. Evolutionary-scale prediction of atomic-level protein structure with a language model. Science, 379(6637):1123–1130, 2023.

Ali Madani, Bryan McCann, Nikhil Naik, Nitish Shirish Keskar, Namrata Anand, Raphael R Eguchi, Po-Ssu Huang, and Richard Socher. Progen: Language modeling for protein generation. arXiv preprint 2004.03497, 2020.

Michele Magrane and UniProt Consortium. Uniprot knowledgebase: a hub of integrated protein data. Database, 2011:bar009, 2011.

Joshua Meier, Roshan Rao, Robert Verkuil, Jason Liu, Tom Sercu, and Alex Rives. Language models enable zero-shot prediction of the effects of mutations on protein function. Advances in neural information processing systems, 34:29287–29303, 2021.

Erik Nijkamp, Jeffrey A Ruffolo, Eli N Weinstein, Nikhil Naik, and Ali Madani. Progen2: exploring the boundaries of protein language models. Cell systems, 14(11):968–978, 2023.

Roshan Rao, Nicholas Bhattacharya, Neil Thomas, Yan Duan, Peter Chen, John Canny, Pieter Abbeel, and Yun Song. Evaluating protein transfer learning with tape. Advances in neural information processing systems, 32, 2019.

Roshan Rao, Joshua Meier, Tom Sercu, Sergey Ovchinnikov, and Alexander Rives. Transformer protein language models are unsupervised structure learners. Biorxiv, pp. 2020–12, 2020.

Jason A. Reuter, Damek V. Spacek, and Michael P. Snyder. High-throughput sequencing technologies. Molecular Cell, pp. 586–597, May 2015. doi: 10.1016/j.molcel.2015.05.004. URL 10.1016/j.molcel.2015.05.004.

Alexander Rives, Joshua Meier, Tom Sercu, Siddharth Goyal, Zeming Lin, Jason Liu, Demi Guo, Myle Ott, C Lawrence Zitnick, Jerry Ma, et al. Biological structure and function emerge from scaling unsupervised learning to 250 million protein sequences. Proceedings of the National Academy of Sciences, 118(15):e2016239118, 2021.

Amir Shanehsazzadeh, David Belanger, and David Dohan. Is transfer learning necessary for protein landscape prediction? arXiv preprint 2011.03443, 2020.

Ilia Shumailov, Zakhar Shumaylov, Yiren Zhao, Nicolas Papernot, Ross Anderson, and Yarin Gal. Ai models collapse when trained on recursively generated data. Nature, 631(8022):755–759, 2024.

Baris E Suzek, Hongzhan Huang, Peter McGarvey, Raja Mazumder, and Cathy H Wu. Uniref: comprehensive and non-redundant uniprot reference clusters. Bioinformatics, 23(10):1282–1288, 2007.

Mihaly Varadi, Stephen Anyango, Mandar Deshpande, Sreenath Nair, Cindy Natassia, Galabina Yordanova, David Yuan, Oana Stroe, Gemma Wood, Agata Laydon, et al. Alphafold protein structure database: massively expanding the structural coverage of protein-sequence space with high-accuracy models. Nucleic acids research, 50(D1):D439–D444, 2022.

Robert Verkuil, Ori Kabeli, Yilun Du, Basile IM Wicky, Lukas F Milles, Justas Dauparas, David Baker, Sergey Ovchinnikov, Tom Sercu, and Alexander Rives. Language models generalize beyond natural proteins. bioRxiv, pp. 2022–12, 2022.

Jesse Vig, Ali Madani, Lav R Varshney, Caiming Xiong, Richard Socher, and Nazneen Fatema Rajani. Bertology meets biology: interpreting attention in protein language models. arXiv preprint 2006.15222, 2020.

Shaojun Wang, Ronghui You, Yunjia Liu, Yi Xiong, and Shanfeng Zhu. Netgo 3.0: protein language model improves large-scale functional annotations. Genomics, Proteomics & Bioinformatics, 21 (2):349–358, 2023.

Minghao Xu, Xinyu Yuan, Santiago Miret, and Jian Tang. Protst: Multi-modality learning of protein sequences and biomedical texts. In International Conference on Machine Learning, pp. 38749– 38767. PMLR, 2023.

Shuwei Yao, Ronghui You, Shaojun Wang, Yi Xiong, Xiaodi Huang, and Shanfeng Zhu. Netgo 2.0: improving large-scale protein function prediction with massive sequence, text, domain, family and network information. Nucleic acids research, 49(W1):W469–W475, 2021.

Ningyu Zhang, Zhen Bi, Xiaozhuan Liang, Siyuan Cheng, Haosen Hong, Shumin Deng, Jiazhang Lian, Qiang Zhang, and Huajun Chen. Ontoprotein: Protein pretraining with gene ontology embedding. arXiv preprint 2201.11147, 2022.

Z Zhang, C Wang, M Xu, V Chenthamarakshan, AC Lozano, P Das, and J Tang. A systematic study of joint representation learning on protein sequences and structures. Preprint at http://arxiv.org/abs/2303.06275, 2023.

Lingyan Zheng, Shuiyang Shi, Pan Fang, Hongning Zhang, Ziqi Pan, Shijie Huang, Weiqi Xia, Honglin Li, Zhenyu Zeng, Shun Zhang, et al. Annopro: an innovative strategy for protein function annotation based on image-like protein representation and multimodal deep learning. bioRxiv, pp. 2023–05, 2023.

Guangjie Zhou, Jun Wang, Xiangliang Zhang, and Guoxian Yu. Deepgoa: predicting gene ontology annotations of proteins via graph convolutional network. In 2019 IEEE International Conference on Bioinformatics and Biomedicine (BIBM), pp. 1836–1841. IEEE, 2019.

